# Variable Gain DNA Nanostructure Charge Amplifiers for Biosensing

**DOI:** 10.1101/2023.08.11.552535

**Authors:** Jacob M. Majikes, Seulki Cho, Thomas E. Cleveland, J. Alexander Liddle, Arvind Balijepalli

**Affiliations:** Microsystems and Nanotechnology Division, National Institute of Standards and Technology, Gaithersburg MD 20899; Biomolecular Measurement Division, National Institute of Standards and Technology, Gaithersburg, MD 20899; Institute for Bioscience and Biotechnology Research, University of Maryland, Rockville, MD 20850

## Abstract

Electronic measurements of engineered nanostructures comprised solely of DNA (DNA nanostructures) enable new signal conditioning modalities in biosensing. Here, we demonstrate how DNA nanostructures that alter their conformation upon binding a nucleic acid analyte drastically, and reversibly, amplify the measured electrochemical signal. This amplification was controlled by the applied electrical field to achieve a response ≈ 2×10^4^ times greater than that measured from DNA hybridization. Because the amplification is independent of the interaction between the analyte and the DNA nanostructure, our approach provides a platform for tuning the response of the system for high performance that is agnostic of the end application. These molecularly precise self-assembled DNA nanostructures when paired with scalable electronic readout can therefore lead the way to highly sensitive multiplexed biosensing.

## Introduction

DNA nanostructures^1^ that can perform local signal conditioning such as amplification have the potential to drastically improve biochemical sensing. DNA nanostructures are fabricated by hybridizing synthetic DNA with a viral scaffold that allows it to ‘fold’ into a predetermined shape. Therefore, these DNA nanostructures offer a modular platform that can be readily functionalized at specific locations with a variety of nanoparticles and biomoleucles,^2^ and can be engineered for dynamic motion with pre-defined rigidity.^3^ To date, a wide array of biosensing proofs-of-concept have utilized these capabilities.^4^ The vast majority of the signals generated by these structures are transduced optically, typically through Förster Resonance Energy Transfer, although several other schemes exist, ranging from gel mobility,^5^ to movement of light-visible nanoparticles,^5^ and to triggered polymerization.^6^ There have also been studies that measure the motion of DNA nanostructures *via* changes in the electrochemical potential, for example the actuation of I-motif using pH,^7^ AC electric field driven motion of DNA nanorods measured optically,^8^ electrochemical measurements of DNA nanostructures,^9^ or capacitance measurements to track nanostructure assembly.^10^

While novel, these demonstrations often fall short of the full potential of DNA nanotechnology in biochemical sensing. Because chemical specificity can be achieved through other means,^11^ the true power of DNA nanostructures lies in their ability to provide local signal conditioning that can drastically improve the signal to noise ratio of the measurement. Here we demonstrate this capability by utilizing DNA nanostructures that alter their conformation^12^ upon binding an analyte. This conformational change displaces a large amount of charge and results in robust amplification of the transduced signal. Furthermore, such an approach can be extended to any analyte type for which an aptamer or DNA binding modification exists potentially allowing parallel multi-analyte sensing.

## Results and Discussion

### Structural Characterization

The DNA nanostructures used in this study take the form of a hinge^12^ comprised of two arms connected by eight short single stranded DNA (ssDNA) tethers as seen in Fig. 1A (*top*). Each arm consists of 20 double stranded DNA (dsDNA) helices arranged in three layers in an 8−4−8 configuration. Each arm has a lock motif placed ≈12 nm from the hinge that allows an analyte to trigger actuation of the structure. The bottom arm is connected to the electrode surface at nine locations using dsDNA tethers or stilts that are 25 nucleotides (≈8.3 nm) long. Representative images from cryogenic electron microscopy (cryo-EM) that were used to validate the structures are shown in Fig. 1A (*bottom left*) with full fields of view in the *Supplementary Information* (SI) section S7. Individual cryo-EM images were combined (SI section S8) to produce a 3D-reconstruction (Fig. 1A; *bottom right*) that validates the design of the DNA nanostructures. Because there is a mismatch in mapping between the desired connections and the relaxed helicity of B-form DNA, enforcing a structure that lies on a square grid result in internal strain and mild torsion as seen in Fig. 1A.

**Fig. 1:**
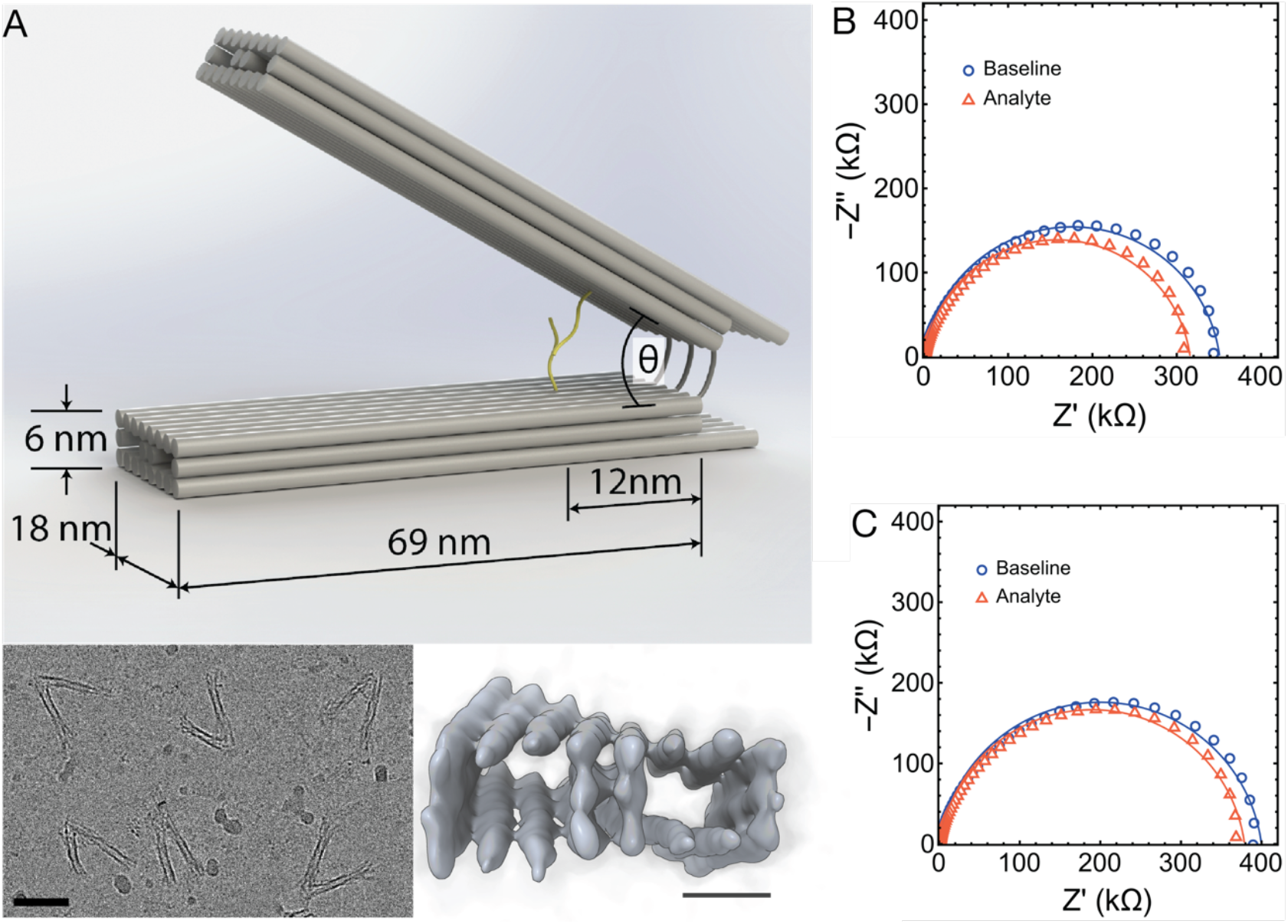
(A) (*top*) Schematic representation of the DNA nanostructures. (*bottom left*) Representative cryogenic electron microscopy (cryo-EM) images of the DNA nanostructures. The scale bar is 50 nm. (*bottom right*) Three-dimensional reconstruction of the structure of the DNA nanostructures from cryo-EM images. The scale bar is 6 nm. (B) and (C) Electrochemical impedance spectroscopy (EIS) of *normally closed* and *normally open* DNA nanostructures in the absence (*blue*) and presence of 1 nmol/L (nM) analyte (*orange*). Solid lines in each case represent fits of a simplified Randles circuit.

The lock motifs (SI section S1) were designed to create two variants, one that remains open (*normally open*) and another that remains closed (*normally closed*) in the absence of analyte. Steric design constraints require the lock motif for the *normally closed* case to be twice the width of the *normally open* structure. Therefore, the *normally closed* structure has a starting angle (in the absence of analyte) of ≈ π/6 rad (30°) and a final angle of ≈ π/2 rad (90°) upon binding an analyte. The *normally open* structure has a starting angle approaching ≈ π/2 rad (90°) and closes more tightly upon exposure to an analyte, to ≈ π/12 rad (15°) due to its shorter lock motif.

### Electrochemical Measurements of Function

We tested the function of the *normally closed* (Fig. 1B) and *normally open* (Fig. 1C) structures by using electrochemical impedance spectroscopy (EIS). In each case, we measured the DNA nanostructures in the absence of analyte (*blue*) and compared them to measurements in the presence of 1 nmol/L (1 nM) analyte (*orange*). By fitting a simplified Randles circuit,^13^ we extracted the circuit parameters for each DNA nanostructure type (parameter table in SI section S2). For both structures we observed in increase in capacitance from a baseline value with no analyte present [*normally closed*, (119±1) nF; *normally open*, (99±1) nF] to a larger value when exposed to 1 nmol/L (1nM) of analyte for 60 minutes [*normally closed*, (143±2) nF; *normally open*, (117±3) nF]. In contrast, the capacitance for ssDNA probe molecules increased by a smaller amount from a baseline value of (38.7±0.3) nF to (41.4±0.6) nF when exposed to 1 nmol/L (1nM) of analyte for 60 minutes (SI section S2). In all cases, the reported error bars were the expanded uncertainty with coverage factor *k*=2 from three independent measurements. To understand the increase in capacitance for both DNA nanostructures we devised a simple model and complementary measurements under an electric field that are discussed next.

### Capacitance Circuit Model

Fig. 2A shows a schematic and corresponding circuit model of the DNA nanostructures that allows us to describe the dependence of *C*_*structure*_ on the angle of the hinge (*θ*) which can be altered by changing the applied DC voltage (*V*_*DC*_). Total capacitance (*C*_*t*_) as a function of *θ* (see SI section S4 for the derivation) was found to be,

**Fig. 2:**
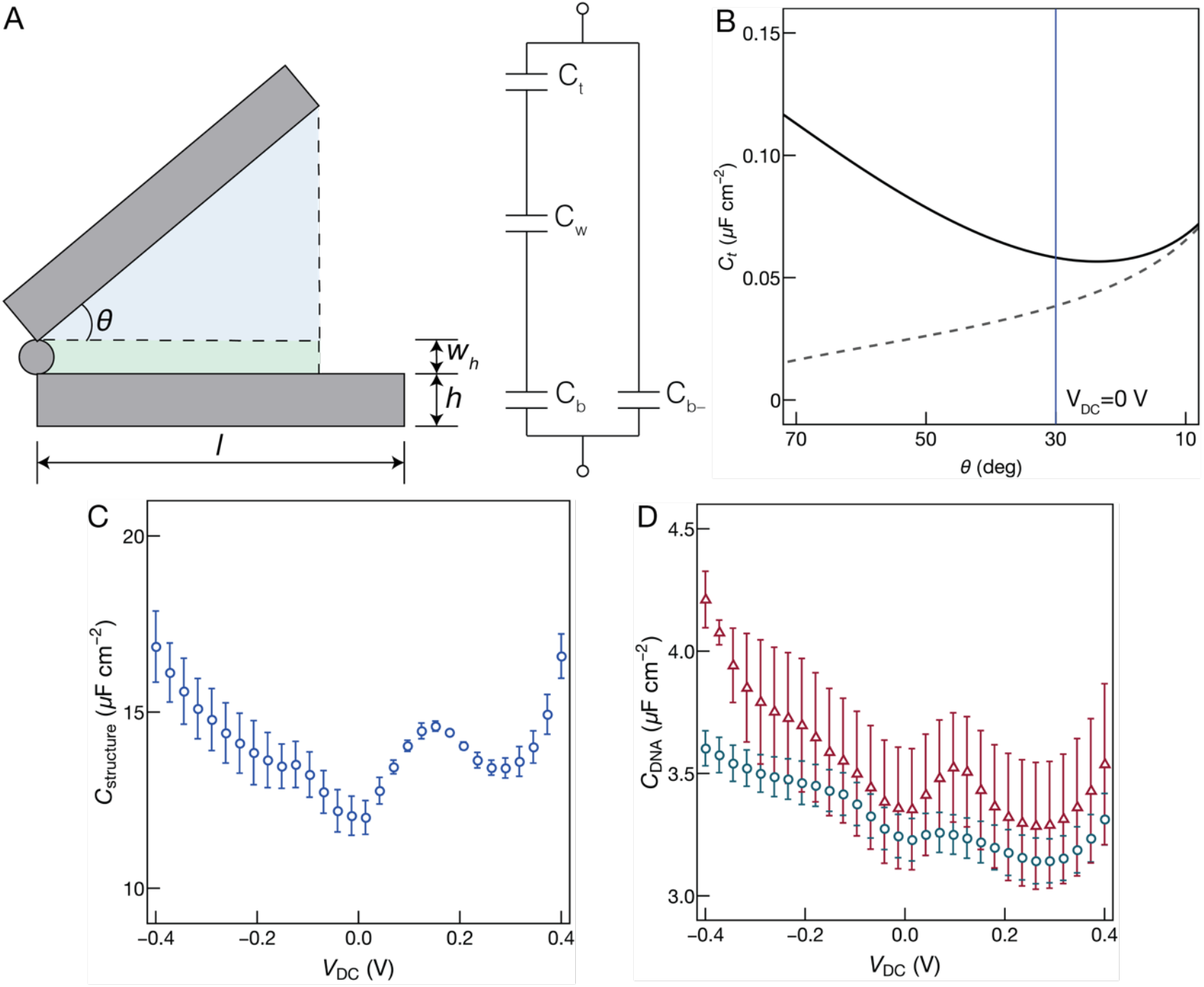
Measurements and modeling of the capacitance of DNA nanostructures as a function of applied DC bias (*V*_*DC*_) relative to an AgCl reference electrode. (A) Schematic and equivalent circuit model of a DNA nanostructures. (B) Total capacitance, *C*_*t*_, as a function of hinge angle, *θ* inferred by solving the equivalent circuit in panel A. (C) Capacitance measurements vs. *V*_*DC*_ of *normally closed* (*blue*) DNA nanostructures (*C*_*structure*_) conjugated to a gold surface in the absence of analyte. The capacitance was measured with an applied AC field with a frequency of 100 Hz and amplitude 20 mV_pk_ summed with *V*_*DC*_. (D) Capacitance measurements vs. *V*_*DC*_ of DNA probe molecules (*C*_*DNA*_) in the absence (*green*) and presence of 1 nM (nmol/L) analyte (*red*).

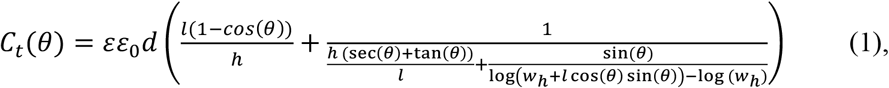

where *l* is the length of the DNA nanostructure, *d* is its depth, *h* is the height of each arm, *w*_*h*_ is the height of the electrolyte layer when the structure is fully closed, *ε* is the dielectric constant and *ε*_*0*_ is the vacuum permittivity. Using the dimensional parameters from Fig. 1 and an estimate of the density of the DNA nanostructures on the electrode surface (SI section S9), we plot *C*_*t*_ as a function of *θ* in Fig. 2B. The *blue* line in the figure shows the expected neutral angle for the *normally closed* structure at *V*_*DC*_=0 V. From Fig. 2 we observe that the expression for *C*_*t*_ qualitatively captures the capacitance behavior as the DNA nanostructure transitions from a closed to an open conformation by the applied electric field. As the structure is closed *C*_*t*_ increases due to the thinning of the electrolyte layer encompassed by the two arms. Conversely, the structure opening results in a broad minimum followed by a slight increase in *C*_*t*_ as *θ* approaches π/2 rad (90°). This increase in *C*_*t*_ is dominated by the capacitance of the bottom arm that is exposed to electrolyte solution as the top arm retracts. The effect of the bottom arm is observed by the dashed line in Fig. 2C, which was calculated by suppressing the second term in Eq. 1. This analysis was validated by measuring the capacitance (*C*_*structure*_) of the *normally closed* case as a function of *V*_*DC*_ in Fig. 2C. The measurements were corrected for the passivation of the electrode surface and the electrical double layer as described in the SI section S3. The measured data show qualitative agreement with the model in Fig. 2B except for the peak centered at *V*_*DC*_ ≈ 0.15 V that are discussed next. Results for the *normally open* case exhibit a similar trend and are shown in the SI section S3.

### Impact of E-Field on dsDNA Spacers

An interesting feature of the plots in Fig. 2C is the presence of the peak centered at *V*_*DC*_ ≈ 0.15 V that is not modeled by Eq. 1. Such behavior has been previously observed when using electric fields to force dsDNA to lie flat on a surface and was found to occur at comparable values of applied voltage for both DNA nanostructures tested in this study.^14,15^ We believe the peak in Fig. 2C is associated with pulling down the nine dsDNA tethers that anchor the structures to the gold surface (Fig. S5.1). To further validate this hypothesis, we hybridized the ssDNA with an analyte and measured *C*_*DNA*_ as a function of *V*_*DC*_ as seen in fig. 2D. While we observed no peak for the ssDNA case (green), we clearly see the emergence of a new peak at ≈0.15 V (*red*) upon hybridizing the ssDNA peaks with a complementary strand. Furthermore, the location of this peak at ≈0.15 V is consistent with the capacitance data in Fig. 2C for the DNA nanostructure strongly indicating that it is related to forcing the nine dsDNA tethers in the structure to lie flat on the electrode surface.

### Capacitance Changes upon Binding Analyte

The change in the capacitance with applied electric field discussed previously can also be initiated by binding a nucleic acid that has a complementary sequence with the lock strand for the *normally open* or *normally closed* structures. While a large change in the measured capacitance upon analyte binding should occur at equilibrium, reaching this state can take a significant amount of time.^16^ This would preclude the structures from use in practical applications such as diagnostics. Here we show that the applied electrical field can be used to accelerate the kinetics of the hinge transition. For both the *normally closed* and *normally open* cases we measured the change in capacitance as a function of *V*_*DC*_ in Fig. 3. In each case the measurements were limited to an incubation time of 15 minutes.

**Fig. 3:**
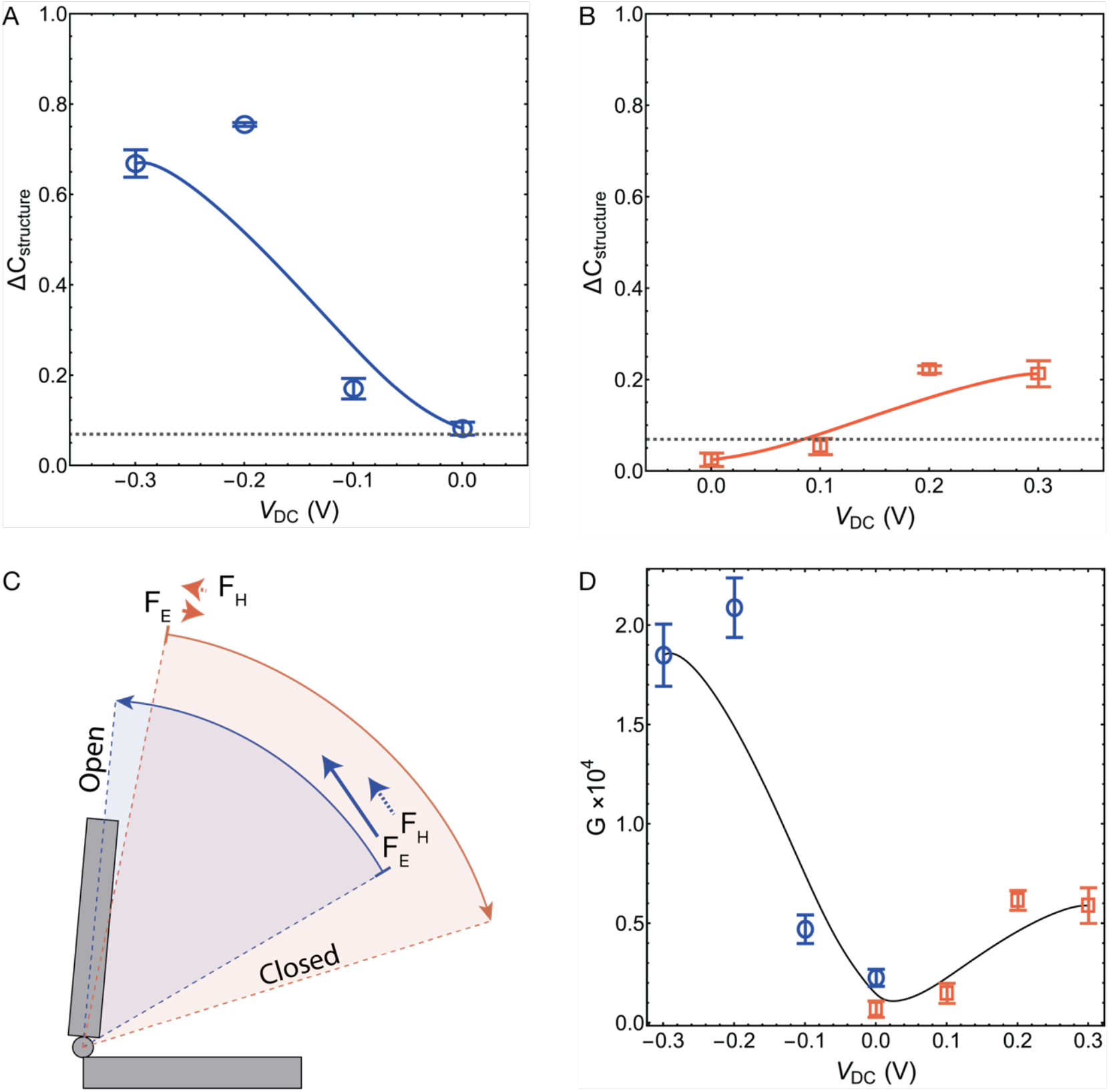
Relative change in capacitance (*ΔC*_*structure*_) and gain of DNA nanostructures relative to the change in capacitance of DNA hybridization (*ΔC*_*ssDNA*_). (A) and (B) Relative change in the capacitance of the *normally closed* and *normally open* DNA nanostructures in the presence of 1 nM (nmol/L) analyte as a function of the DC bias (*V*_*DC*_) relative to the absence of analyte. The dashed lines in each panel show *ΔC*_*ssDNA*_ in the presence of 1 nM (nmol/L) of DNA analyte. (C) Schematic of the forces from the hinge spring constant, F_H_, and the portion of electric force normal to the hinge, F_E_, at the relevant angles for the two structures as described in the main text. (D) Gain (*G*) of the *normally closed* (*blue*) and *normally open* (*orange*) DNA nanostructures over the DNA hybridization case. In all the plots, the error bars in the measurements represent the expanded uncertainty with coverage factor *k*=2.

Fig. 3A shows the relative change in the DNA nanostructures capacitance (*ΔC*_*structure*_) for the *normally closed* case before and after incubation with 1 nmol/L (nM) of DNA of a complementary sequence to the lock strand. Increasing the negative voltage forces the structures to open, upon which we see an increase in *ΔC*_*structure*_that saturates at Δ −0.3 V. Upon flushing the fluidic cell with RBS to remove excess analyte, we observed no further change in *ΔC*_*structure*_ indicating that the analyte binding the lock strands prevented the structures from reverting to a closed state. On average, we observed that *ΔC*_*structure*_ changed by 0.08±0.01, 0.17±0.02, 0.8±0.2 and 0.7±0.3 for an applied *V*_*DC*_ of 0 V, −0.1 V, −0.2 V and −0.3 V respectively for the *normally closed* cases. The observed values of *ΔC*_*structure*_ represent an ≈11-fold increase compared with the hybridization of ssDNA probe strands with a complementary sequence (*ΔC*_*ssDNA*_=0.07±0.01). All error bars represent the expanded uncertainty with coverage factor *k*=2. A smaller enhancement in *ΔC*_*structure*_ was observed for the *normally open* case as seen in Fig. 3B. Here *ΔC*_*structure*_ was found to be 0.01±0.01, 0.05±0.02, 0.2±0.08 and 0.2±0.02 for an applied *V*_*DC*_ of 0 V, +0.1 V, +0.2 V and +0.3 V respectively. While this represented an ≈3-fold higher improvement over the ssDNA hybridization case, it was comparatively lower than the *normally closed* results.

As seen in Fig. 3C, the difference in the enhancement between the two structures can be attributed to the effectiveness of the electric field (*E*) at actuating the hinge structure. When the hinge is closed (*θ →* 0 rad or 0°), *E* is orthogonal to the top arm resulting in a large electric force (*F*_*E*_) that works with the hinge restoring force (*F*_*H*_) to efficiently open the structure. When the hinge is open (*θ* → π/2 rad or 90°), *E* is parallel to the top arm and therefore applies a negligible *F*_*E*_. At the open position, the hinge spring is at its energy minimum also resulting in a negligible *F*_*H*_. In short, *F*_*E*_ is more effective at opening this structure than closing it. Importantly, this subtle difference translates into a drastically different signal gain.

### Signal Amplification upon Binding Analyte

The signal gain (*G*) relative to the ssDNA case (Fig. 3D) was estimated by scaling *ΔC*_*structure*_ and *ΔC*_*ssDNA*_ with the ratio of the densities of DNA probe molecules (*ρ*_*ssDNA*_) attached to the electrode surface to the densities of the DNA nanostructures (*ρ*_*structure*_) and given by,

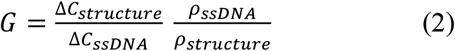

The values of *ρ*_*structure*_ and *ρ*_*ssDNA*_ were measured chronocoulometrically^17^ (SI section S9) to be (6.21±0.004)×10^8^ cm^-2^ and (1.19±0.001)×10^12^ cm^-2^ respectively. From Fig. 3D, *G* increases monotonically with negative voltages for the *normally closed* structures and with positive voltages for the *normally open* structures. This trend mirrors the change in the measured capacitance in Fig. 3A and 3B. The estimated gain peaks at (2.1±0.1)×10^4^ for the *normally closed* structures and (0.6±0.05)×10^4^ for the *normally open* case before reaching a plateau. This maximum indicates that the structures are strongly responsive to *E*, requiring low voltages to be driven to change their conformations. This significant signal amplification can be further improved by structure modification to increase mobile charge, by more effectively utilizing the hinge restoring force, *F*_*H*_, and by more closely binding the hinge to the surface, i.e., removing the anchor stilts.

## Conclusions

As DNA nanostructures are readily designed into custom geometries, and increasingly support engineered dynamic properties, they have great potential for signal conditioning in electric measurements of biomolecular binding. As we show here, simple hinge-based geometries can be adapted for dramatically improved gain as high as ≈ 2×10^4^ compared to ssDNA hybridization and be used reversibly for multiple measurements. Beyond the pathways to improved signal amplification these systems also have the potential to modify the energetics and kinetics of binding for arbitrary analytes. This will open broad research directions to develop frameworks to incorporate different lock types such as aptamers, and for modifying the dynamic motion of the structures.

## Materials and Methods

### DNA Nanostructure synthesis

The sequences used for the DNA nanostructures were identical previous work,^12^ except for the modifications discussed in SI section S1. The P8064 M13 sequence was used as a scaffold at a concentration of 35 nmol/L. The staples, suspended in 40 mmol/L Tris, 20 mmol/L acetic acid, 1 mmol/L EDTA (TAE) buffer and 12.5 mmol/L MgCl_2_ in deionized water (DIW), were introduced at a 10-fold higher concentration than the scaffold (350 nmol/L). The DNA staples were annealed with the scaffold for 12 hours from 80 °C to 4 °C. A centrifugal spin filter with a 1.67×10^-19^ g (100 kDa) molecular weight cutoff was used to remove excess staples through five iterative additions of 0.5 mL of buffer. The samples were stored at 4 °C. DNA samples for Cryo-EM microscopy were annealed with 80 nmol/L scaffold.

### Gold Electrode Functionalization

Commercially sourced gold electrodes were prepared for electrochemical measurements by cleaning and modifying them with thiol chemistry. The electrodes were first cleaned by immersing them in nanostrip for 10 minutes followed by thoroughly rinsing them in DIW. The electrodes were then further cleaned electrochemically in a solution of 50 mmol/L of sulfuric acid (H_2_SO_4_) by sweeping a voltage from −0.5 V to +1 V relative to an Ag/AgCl pseudo reference electrode at a rate of 0.1 V/s. The sweeps were repeated for a minimum of 10 cycles and until the reduction peak reached a steady state. The substrates were then washed with DIW. A running buffer solution (RBS) comprised of TAE with 50 mmol/L NaCl, 12.5 mmol/L MgCl_2_, 200 µmol/L K_3_ Fe(CN)_6_ and 200 µmol/L K_4_ Fe(CN)_6_ was prepared and used both for modifying the surfaces and for the electrochemical measurements. The cleaned chips were incubated for >18 hours at room temperature in RBS with 10 nmol/L of the DNA nanostructures, 10 nmol/L of mercaptohexanol (MCH) for surface passivation of the unreacted gold and 10 µmol/L Tris (2-carboxyethyl) phosphine (TCEP) as reducing agent. The chips were then thoroughly washed with RBS before being used for electrochemical measurements. Chronocoulometric measurements were realized by adding 50 µmol/L of Hexaammineruthenium(III) chloride (RuHex) to RBS.

### Electrochemical Measurements

All electrochemical impedance spectroscopy measurements were performed with a two-electrode electrochemical cell with a 200 µL sample volume. An impedance analyzer was used to apply an AC voltage with an amplitude of 20 mV_pk_ over a frequency range of 0.5 Hz to 200 kHz and measure the resulting current. Baseline measurements were performed with RBS and again after incubating with analyte for an incubation time specified in the main text followed by rinsing the cell with RBS to remove excess analyte. All capacitance measurements were performed at a fixed frequency of 100 Hz, AC voltage amplitude of 20 mV_pk_ and by sweeping the DC voltage from −0.4 V to +0.4 V relative to an Ag/AgCl pseudo reference electrode.

### Cryogenic Electron Microscopy

CryoEM measurements were performed to reconstruct the DNA nanostructures. The large size of the structures complicated the analysis as did the symmetry of the arms. However, we were able to leverage the symmetry of the structures to perform the reconstruction by not distinguishing between the each arm. A detailed description of the measurements and reconstruction techniques are provided in SI section S8.

## Notes

### Competing Interest Statement

The authors have declared no competing interest.

