## SupplementaryInformation for "Variable Gain DNA Nanostructure Charge Amplifiers for Biosensing"

### S1: DNA Nanostructure Size and Design

The DNA nanostructure, as discussed in the main text, was designed to be approximately ≈ 69 nm long, ≈ 6 nm tall, and ≈ 18 nm wide. The structure used here is consistent with the one labeled in the cited work[1] as nDFS.B. The lock motif was modified, as were the 9 positions which anchor the structure to the electrode.

It should be noted that the hinge design is on the square lattice (helices are connected at π/2 rad (90 ˚) angles. As dsDNA is only relaxed at a helicity of 10.5 nucleotides per full rotation, and connection points can only occur between helices at integer numbers of nucleotides, 3D structures built in this way are inherently strained. As seen in Fig. 1 of the main text, this strain is compensated by applying a slight torque on the hinge arms.

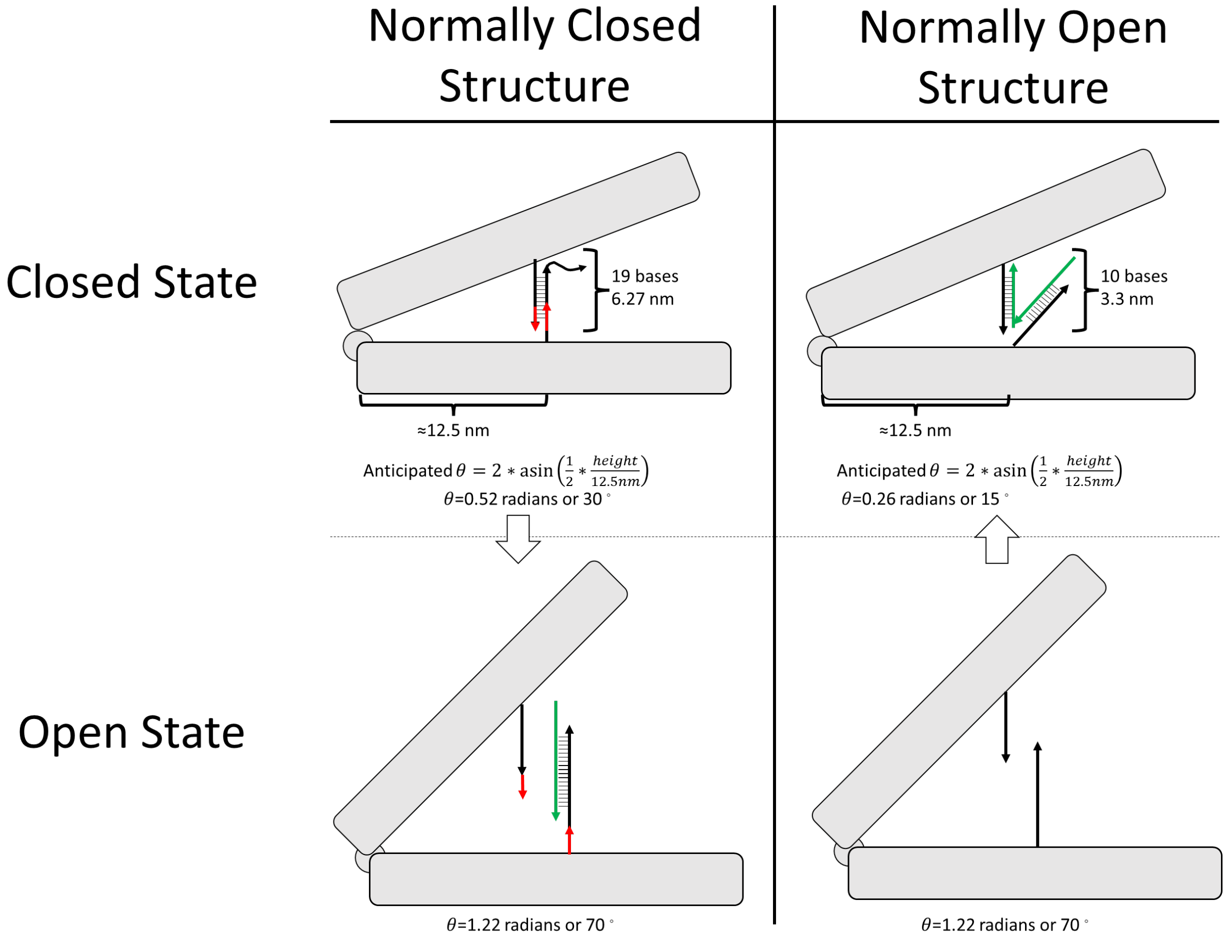

Fig. S1.1: Schematics of the normally open and normally closed structures, the hybridization of their lock motifs in their open and closed states, and the estimated angle those lock motifs should enforce.

Relevant strand sequences

| | |
|---|---|
| Base Analyte | TTATGTGACCGACGAGACTA |
| Analyte_W/Toe | AGTTC TTATGTGACCGACGAGACTA |
| New Analyte Complement | TAGTCTCGTCGGTCACATAA |

| | |
|---|---|
| Thiolated Strand | /5DTPA/TTTACCGGAAGCAAACTGCTTCAAAGCG |
| Unmodified Top Lock staple | ATAAGCGGAATTATCATCATATTTTAAATACCGTTC |
| Unmodified Bottom Lock staple | AAGATGATGAAACAAATCAATATAAGAATCCTTT |

Where Analyte_W/Toe was used to allow for preliminary strand exchange experiments. The Unmodified Top Lock staple and Unmodified Bottom Lock staple sequences correspond to CSBottom and CSTop.[1]
These sequences were used to generate the lock motif sequences, given below.

Red indicates the toehold sequence for strand displacement, green indicates spacer bases, Blue indicates the respective base sequence.

| Default Closed Structure | | Default Open Structure | |
|---|---|---|---|
| Sticky Seq. Top | TTGACCGACGAGACTAGTG | Sticky Seq. Top | TAGTCTCGTC |
| Sticky Seq. Bottom | TCACTAGTCTCGTCGGTCACATAA | Sticky Seq. Bottom | GGTCACATAA |
| Closed Top | ATAAGCGGAATTATCATCATATTTTAAATACCG TTCTTGACCGACGAGACTAGTG | Open Top | ATAAGCGGAATTATCATCATATTTTA AATACCGTTCTAGTCTCGTC |
| Closed Bottom | AAGATGATGAAACAAATCAATATAAGAATCCTT TTCACTAGTCTCGTCGGTCACATAA | Open Bottom | AAGATGATGAAACAAATCAATATAAG AATCCTTTGGTCACATAA |

The anchor strands used in the cited work were modified to remove the ssDNA spacer between the dsDNA stilts and the DNA nanostructure.

| Name | Sequence |
|---|---|
| Surface1 | TTCAGAGGCAGGAAACAAAAAATAACGGCTTAATTGCGCTTTGAAGCAGTTTGCTTCCGGT |
| Surface2 | GTTTTATAACTAACAAAGAAAGAAACAAGGTAATTGCGCTTTGAAGCAGTTTGCTTCCGGT |
| Surface3 | AGAATCGCATCTTACCAACGCTAATTGAAGCCCGCTTTGAAGCAGTTTGCTTCCGGT |
| Surface4 | TATCATATTATTATTTATCCCAATAAGGCTTACGCTTTGAAGCAGTTTGCTTCCGGT |
| Surface5 | AGCGCTAACCTTTACAGAGAGAATAAGCCGTTCGCTTTGAAGCAGTTTGCTTCCGGT |
| Surface6 | TATTACGCAATACCGACCGTGTGACTGTTTAGCGCTTTGAAGCAGTTTGCTTCCGGT |
| Surface7 | GCATTTTCGGTCATAGTCAGAGCCGCCAAACGAAAAGACCCGCTTTGAAGCAGTTTGCTTCCGGT |
| Surface8 | CAGTAGCGCATATGGTTTACCAGCAGACTCCTCGCTTTGAAGCAGTTTGCTTCCGGT |
| Surface9 | GCACCATTATATTGACGGAAATTAGTTACCAGCGCTTTGAAGCAGTTTGCTTCCGGT |
| Thiol Anchor | /5DTPA/TTTACCGGAAGCAAACTGCTTCAAAGCG |

### S2: Electrochemical Measurements Methods and Measurements

Electrochemical impedance spectroscopy (EIS) measurements modeled using a Randles circuit[2] (Fig. S2.1A) that represents the solution resistance (*Rs*), the resistance of the electrode interface (*Rp*) and a constant phase element representing the capacitance of the interface (*Cp*). were performed for ssDNA (Fig. S2.1B), and the *normally closed* (Fig. S2.1C) and *normally open* (Fig. S2.1D) DNA nanostructures. A complete table of the fit parameters for the model in Fig. S2.1A are shown in Table S2.1.

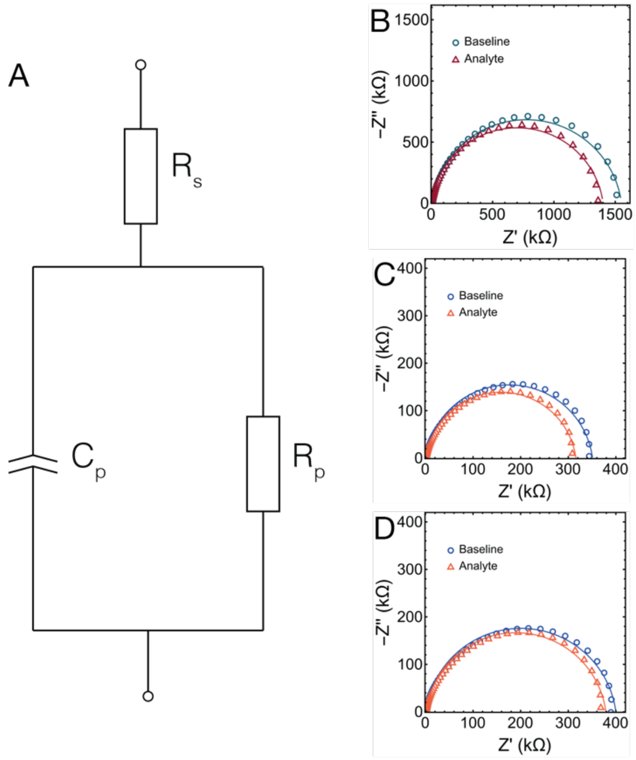

Fig. S2.1: (A) Equivalent circuit diagram used to model electrochemical impedance spectroscopy (EIS) measurements. (B) EIS measurements of ssDNA (*green*) and upon adding 1 nmol/L of complementary analyte (*red*). (C) EIS measurements of the *normally closed* DNA nanostructures (*blue*) and upon adding 1 nmol/L of complementary analyte (*orange*). (D) EIS measurements of the *normally open* DNA nanostructures (*blue*) and upon adding 1 nmol/L of complementary analyte (*orange*).

**Table S2.1: Fit parameters for the circuit in Fig. S2.1**

| | **Rs (Ω)** | **Rp (kΩ)** | **Cp (nF)** | **n** |
|---|---|---|---|---|
| **ssDNA Baseline** | 918±5 | 1551±12 | 38.7±0.3 | 0.89±0.00 |
| **ssDNA Analyte (1 nmol/L)** | 948±0.3 | 1360±17 | 41.4±0.6 | 0.90±0.00 |
| **Normally Closed DNA Nanostructure Baseline** | 898±2 | 335±4 | 119±1 | 0.92±0.00 |
| **Normally Closed DNA Nanostructure Analyte (1 nmol/L)** | 726±2 | 307±1 | 143±2 | 0.91±0.00 |
| **Normally Open DNA Nanostructure Baseline** | 936±1 | 375±7 | 99±1 | 0.92±0.00 |
| **Normally Open DNA Nanostructure Analyte (1 nmol/L)** | 963±1 | 358±7 | 117±3 | 0.91±0.00 |

### S3: Electrical Double Layer and Electrode Passivation Corrections to DNA nanostructure Capacitance

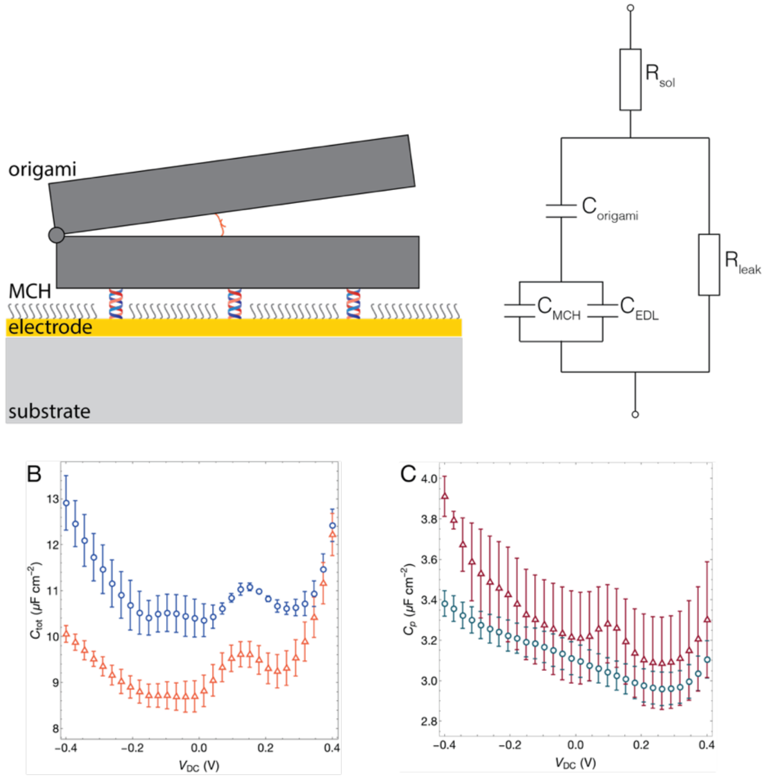

Fig S3.1: (A) (*left*) Schematic representation of the DNA nanostructures anchored to a gold electrode with the unreacted gold surface passivated with 6-mercapto-1-hexanol (MCH) and (*right*) the equivalent circuit model for this system. (B) Capacitance measurements of the *normally closed* (*blue*) and *normally open* (*orange*) cases as a function of a DC bias potential ($V_{DC}$). (C) Capacitance measurements of single stranded DNA probes on a gold surface (*green*) and upon adding 1 nmol/L of analyte with a complementary sequence to the probes (*red*) as a function of $V_{DC}$.

Fig. S3.1A (*left*) shows a schematic of the DNA nanostructures measurement on a gold electrode. Each DNA nanostructure structure is attached to the gold surface with 9 double stranded DNA (dsDNA) strands and the unreacted gold surface is passivated with 6-mercapto-1-hexanol (MCH). The system is modeled with an equivalent electrochemical circuit[3] shown in Fig. S3.1A (*right*). The effective capacitance ($C_{tot}$) of the system can be separated into its constituent elements shown in the figure and represented by the equation,

$$\frac{1}{C_{tot}} = \frac{1}{C_{structure}} + \frac{1}{C_m}, \qquad \text{(S3.1)}$$

where $C_{structure}$ is the capacitance of the DNA nanostructures and $C_m$=$C_{MCH}$+$C_{EDL}$ is the effective capacitance of the passivation surface that includes the capacitance of the MCH layer ($C_{MCH}$) and that of the electrical double layer ($C_{EDL}$) of unpassivated electrode surface.[4,5] We estimate the effective value of $C_m$ as a function of the applied DC bias relative to an AgCl reference electrode ($V_{DC}$) by measuring a surface functionalized with MCH but without the DNA nanostructures. By using Eq. S3.1, we then isolate the contribution of $C_{structure}$ from the overall capacitance in Fig. S3.

**S4. DNA Nanostructure Capacitance Model as a Function of Opening Angle**

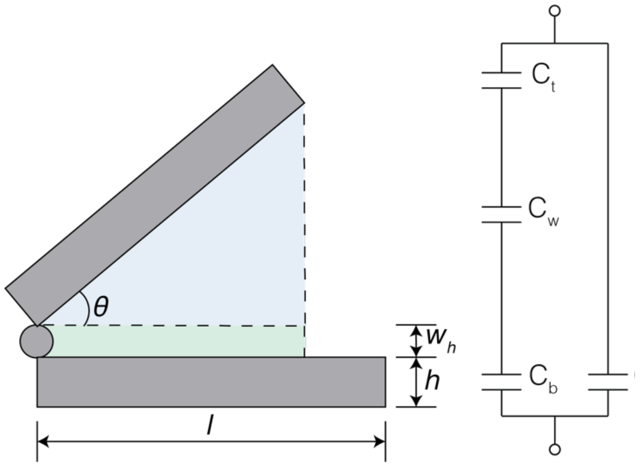

Fig. S4.1: Schematic and equivalent circuit model of a DNA nanostructures

To model the capacitance of the DNA nanostructure as a function of hinge angle, the capacitance was divided into four components, (i) the bottom arm of the DNA nanostructure,

$$C_b = \frac{\varepsilon\varepsilon_0 l\ d\ cos(\theta)}{h}, \quad \text{(S4.1)}$$

(ii) the water layer between the two arms of the DNA nanostructure,

$$C_w = \varepsilon\varepsilon_0 \int_0^{l\cos(\theta)} \frac{d}{h + x\sin(\theta)} dx = \varepsilon\varepsilon_0 d \csc(\theta)(\ln(w_h + l\cos(\theta)\sin(\theta)) - \ln(w_h)), \quad \text{(S4.2)}$$

(iii) the top arm of the DNA nanostructure,

$$C_t = \varepsilon\varepsilon_0 \frac{l\cos(\theta)}{h\sin(\theta)}, \quad \text{(S4.3)}$$

and (iv) the portion of the bottom arm that is exposed to solution as the top arm actuates and ignoring the contribution from the electrical double layer,

$$C_{b^-} = \varepsilon\varepsilon_0 \frac{l\ d(1-\cos(\theta))}{h}, \quad \text{(S4.4)}$$

where $\varepsilon_0$ is vacuum permittivity, $\varepsilon$ is the dielectric constant, $l$ is the hinge length, $d$ is the hinge depth, $h$ is the height of each hinge, $w_h$ is the length of the spacer bases which connect the hinge halves, $\theta$ is the angle of the hinge, $\varepsilon$ is the dielectric constant and $\varepsilon_0$ is Faraday's constant. The dielectric constant of the DNA nanostructures is assumed to be equal to that of the buffer solution since the structure is liquid filled.

Eq. S4.1 – S4.2 can be combined to obtain the final expression in Eq. 1 by using,

$$C_t(\theta) = \left(\frac{1}{C_b} + \frac{1}{C_w} + \frac{1}{C_t}\right)^{-1} + C_{b^-} \quad \text{(S4.5)}$$

**S5: Hypothesized Tether or Stilt Behavior**

The original hinge origami design utilized 9 identical sticky end extensions which would hybridize to 9 copies of a thiolated sequence. In the cited work there was a large ssDNA spacer on these stilts which we removed after preliminary results indicating that this obscured the capacitance data by allowing the structure to be too mobile.

For the nanostructures in this manuscript the 9 anchor positions were comprised of 25 nucleotides of dsDNA, approximately 8.3 nm in length. These could be considered freely jointed at both the gold surface and the origami, and as such, could be expected to collapse as shown in fig. S5.1. The similarity in the peak in capacitance as a function of DC bias between the hinge nanostructure and plain dsDNA would support that at 0.15V the dsDNA is forced to lie flat on the electrode surface.

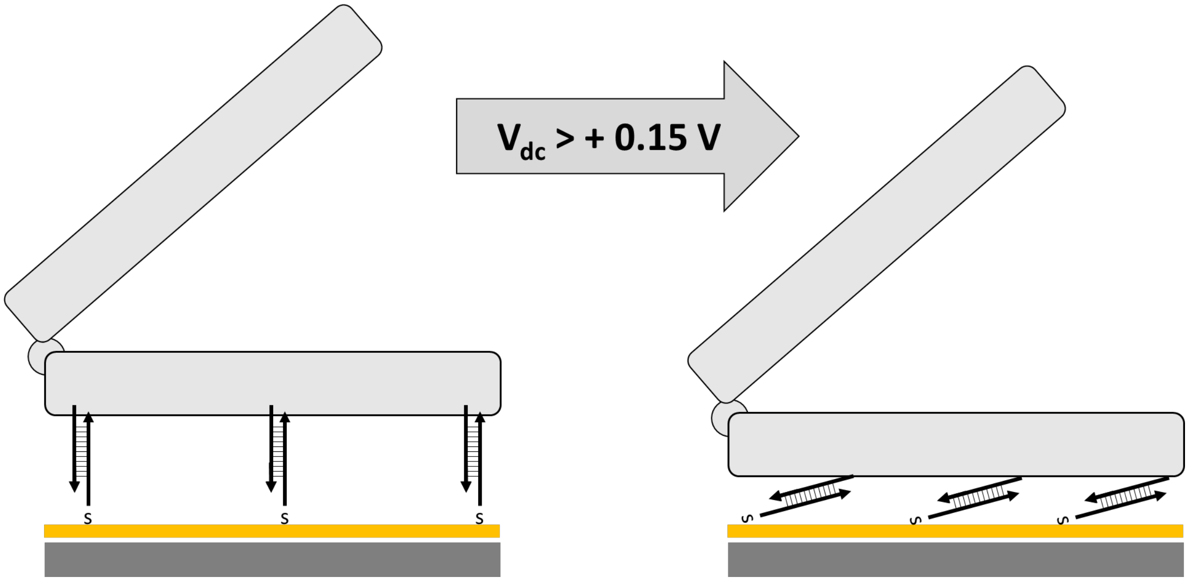

Fig. S5.1: Schematic of the 25 base anchor strands, and how they could be forced to lie collectively flat on the surface of the electrode by an applied positive voltage.

### S6: AFM images of Hinge DNA Nanostructures

AFM was found to be an unsatisfying method to characterize the hinge structure, as under typical conditions of liquid imaging mica in 12.5 mM $Mg^{2+}$ the structures were sufficiently tall and sufficiently weakly bound o the surface to result in poor image quality.

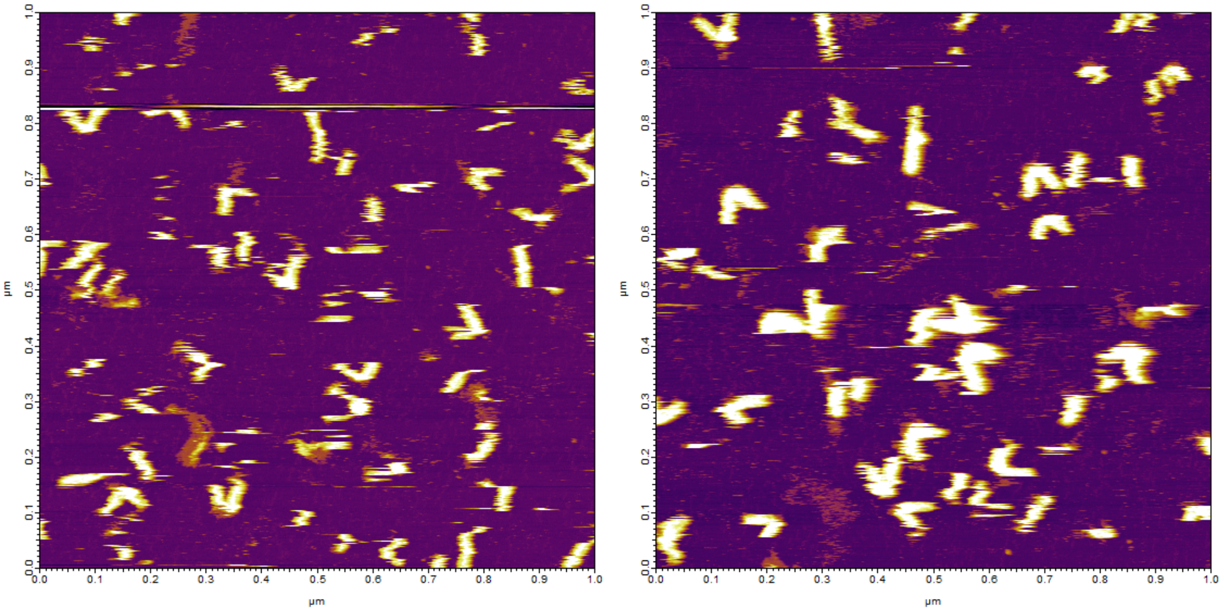

Fig. S6.1: AFM images of the normally closed hinge nanostructure imaged on mica

**S7: Cryo-EM Images of Hinge DNA Nanostructures**
Below are representative examples of Cryo-EM micrographs of the normally closed hinge nanostructure.

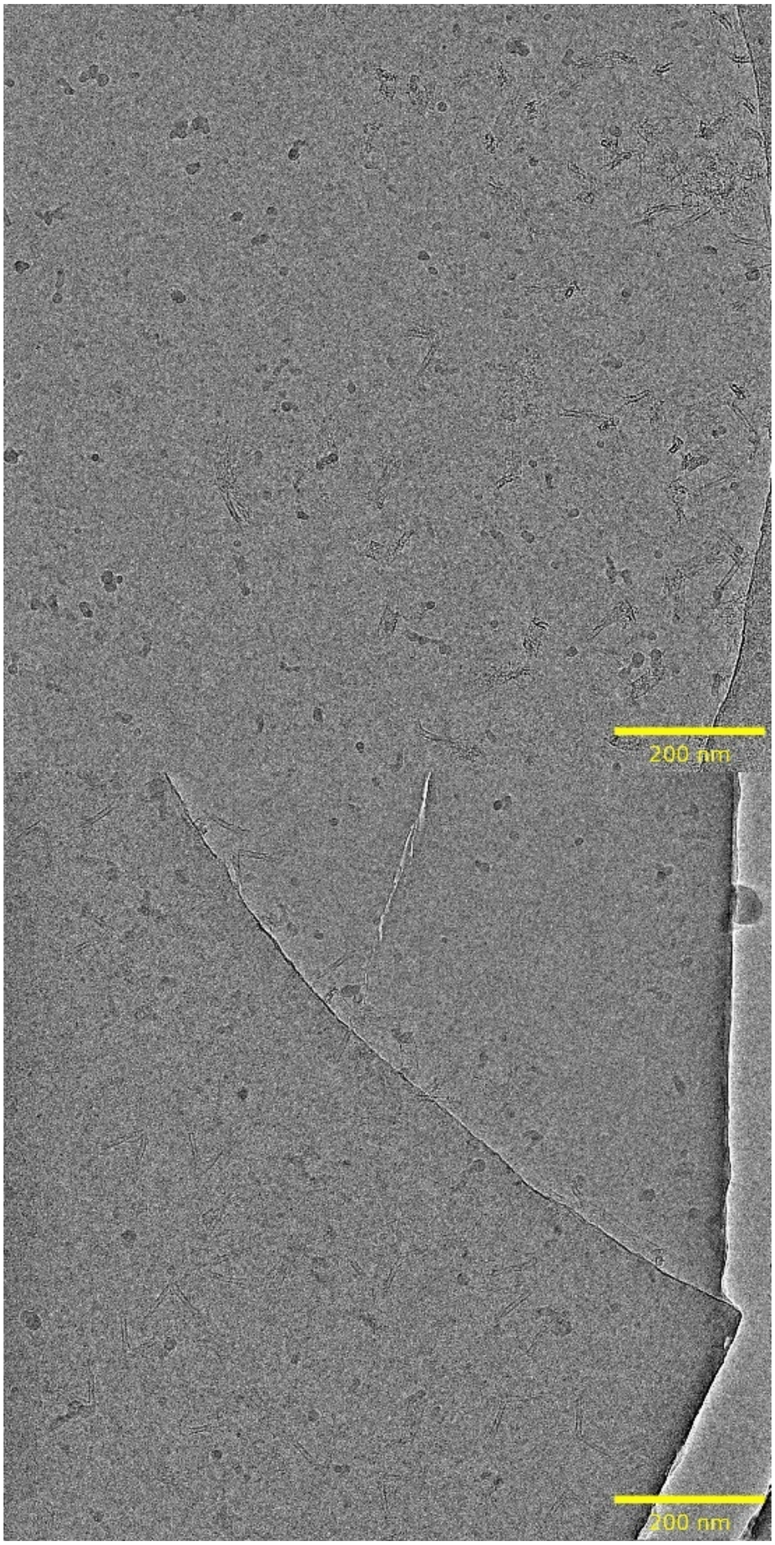

Fig. S7.1: Representative micrographs of the hinge nanostructure

### S8: Cryo-EM Methods and 3D Reconstruction

Cryogenic Electron Microscopy measurements were performed using R3.5/1 micromachined holey carbon with 200 mesh copper supports (Quantifoil*) and were glow discharged using a Pelco easiGlow system (25 mA, 25 s glow time, 10 s hold time) prior to sample application. 3 uL of DNA nanostructure solution was applied to the grid by pipetting, then blotted with filter paper for 5.5 s and plunge-frozen into liquid ethane using a Vitrobot (Thermo Fisher Scientific/"TFS"). Microscopy was performed on a Glacios transmission electron microscope (TFS) operated at 200 kV, using a Falcon 4i direct electron detector (TFS). SerialEM[6] was used for automated data collection. Micrographs were collected in Electron Event Representation (EER) mode at a nominal magnification of 57,000x, corresponding to a physical pixel size of 2.49 Å, with an exposure time of 24.98 s and a total dose of 24.8 e/Å$^2$. Motion correction was performed using Relion v3.1.2[7] with 4K pixel rendering and EER fractionation of 316 frames, corresponding to a dose of 0.987 e/Å$^2$ per fraction. Roughly 450 particles (hinge halves) were picked manually from a subset of micrographs and extracted using a box size of 200 pixels (498 Å). Extracted particles were subjected to 2D classification in Relion to generate templates, which were then used for automated picking, extraction, and 2D classification of particles from all micrographs. Of the 6,687 total particles, 4,002 particles belonged to classes showing high-resolution features (i.e. DNA helices) and were selected for initial model generation and 3D auto-refinement in Relion. For 2D classification and 3D refinement steps, circular masks of 400 Å were applied, effectively restricting refinement to the central portion of the hinge arm to visualize the arrangement of helices. Masking and postprocessing procedures were then performed within Relion, yielding a final map with a reported resolution of 20 Å.

The 2D class images of the hinge arm prior to 3D reconstruction are given below in Fig. 8.1.

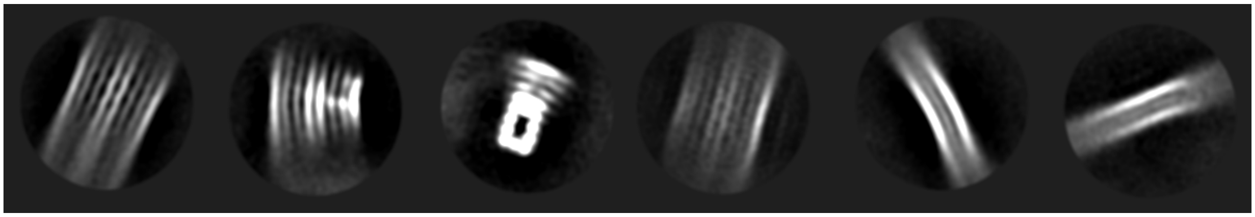

Fig. S8.1: 2D class montage of the hinge arms used to create the 3D reconstruction given in Fig. 1 of the main text.

*Certain commercial equipment, instruments, or materials are identified in this paper to specify the experimental procedure adequately. Such identifications are not intended to imply recommendation or endorsement by the National Institute of Standards and Technology, nor it is intended to imply that the materials or equipment identified are necessarily the best available for the purpose.

### S9: Chronocoulometric Measurements to Estimate Surface Density

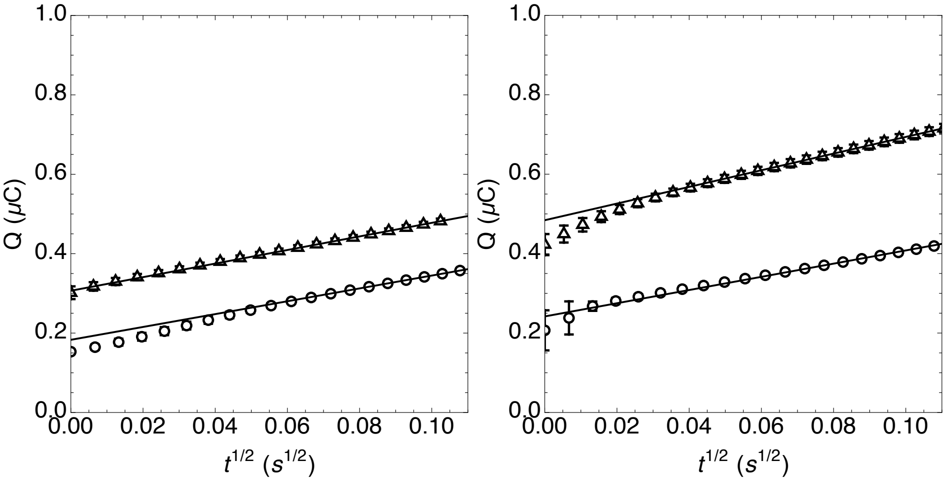

Fig. S9.1: Chronocoulometric response for hybridized dsDNA (*left*) and DNA nanostructure (*right*) in the presence (*triangles*) and absence (*circles*) of 50 µmol/L (µM) of Hexaammineruthenium(III) chloride (RuHex). The solid lines indicate the fit to the data that were used to determine the x-intercept.

Chronocoulometric measurments were used to estimate the surface concentration of both the hybridized dsDNA and the DNA nanostructures following methods in the literature.[8] Fig. S9.1 shows the response curves for each case that was used to estimate the surface concentration[2,8] of each species using the integrated Cotrell equation,

$$Q = \frac{2n\,FAD_0{}^{1/2}C_0}{\pi^{1/2}}\,t^{1/2} + Q_{dl} + n\,FA\,\Gamma_0, \qquad \text{(S9.1)}$$

where $n$ is the number of electrons per molecule, $F$ is the Faraday constant, $A$ is the electrode area, $D_0$ is the diffusion constant, $C_0$ is the bulk concentration, $Q_{dl}$ is the capacitive charge and $nFA\Gamma_0$ is the charge from the reduction of the adsorbed redox marker. The x-intercept of the lines in Fig. S9.1 include the charge from the electric double layer and the adsorbed redox potential, the difference between the case with and without the redox marker is used to estimate the surface excess of adsorbed marker.
